# CPTAC Pancancer Phosphoproteomics Kinase Enrichment Analysis with ProKAP Provides Insights into Immunogenic Signaling Pathways

**DOI:** 10.1101/2021.11.05.450069

**Authors:** Anna Calinawan, Weiping Ma, John Erol Evangelista, Boris Reva, Francesca Petralia, Avi Ma’ayan, Pei Wang

**Affiliations:** Department of Genetics and Genomic Sciences, Icahn School of Medicine at Mount Sinai, New York, NY, 10029, USA; Department of Pharmacological Sciences, Mount Sinai Center for Bioinformatics, Icahn School of Medicine at Mount Sinai, New York, NY 10029

**Keywords:** cancer proteomics, data visualization, kinase enrichment analysis, gene set enrichment analysis, patient cohorts, appyters

## Abstract

The National Cancer Institute (NCI) Clinical Proteomic Tumor Analysis Consortium (CPTAC) initiative has generated extensive phosphoproteomics and proteomics data for tumor and tumor-adjacent normal tissue across multiple cancer types. This dataset provides an unprecedented opportunity to systematically characterize pan-cancer kinase activities, which is essential for coupling tumor subtypes with kinase inhibitors as potential treatment. In this work, we performed Kinase Enrichment Analysis (KEA) using a CPTAC phosphoproteomics dataset to identify putative differences in kinase state between tumor and normal tissues within and across five types of cancer. We then implemented an interactive web-portal, the ProTrack Kinase Activity Portal (ProKAP), for querying, visualizing, and downloading the derived pan-cancer kinase activity scores together with the corresponding sample metadata, and protein and phosphoprotein expression profiles. To illustrate the usage of this digital resource, we analyzed the association between kinase activity scores and immune subtypes of clear cell renal cell carcinoma (ccRCC) derived from the CPTAC ccRCC study. We found multiple kinases, whose inhibition has been suggested to have therapeutic effect in other tumor types, are highly active in CD8+-enriched ccRCC tumors. The ProTrack Kinase Activity Portal (ProKAP) is available at: https://pancan-kea3.cptac-data-view.org.

## Introduction

Protein kinases are key regulators of signal transduction pathways by their function as reversible switches. By phosphorylating the threonine, serine, or tyrosine residues of other proteins, including other protein kinases and phosphatases, kinases modulate the substrate activity, intracellular localization, and half-life [1]. Notably, protein kinases are therapeutically actionable since there are many small molecules and approved drugs that target the human kinome. Dysregulation of phosphorylation-driven signal transduction plays a central role in many cancers, and protein kinases have been demonstrated as effective targets for many cancer treatments. For example, tyrosine kinase inhibitors have improved the outcome of several cancer types, including renal cell carcinoma [2], gastrointestinal stromal tumors (GISTs) [3], and hepatocellular carcinoma [4]. Mapping the best kinase inhibitors for a tumor type for an individual patient is not trivial. To develop a rational approach for determining the kinase inhibitors that will be most effective in blocking the proliferation and survival of tumors, large scale omics profiling can be used to identify dysregulated proteins and phosphoproteins differentiating tumor and normal tissues [5-7]. One earlier resource for such data is the Cancer Genome Atlas (TCGA) [8] which provides comprehensive multi-omics profiling of tumors from thousands of patients including the results from reverse-phase protein arrays (RPPA) [9]. While RPPA provides informative abundance measurements for hundreds of proteins and phosphoproteins, it does not impart comprehensive coverage of the tumor proteome and phosphoproteome at a global scale. To address this gap, the National Cancer Institute (NCI) established the Clinical Proteomic Tumor Analysis Consortium (CPTAC) program which utilizes mass spectrometry proteomics to deeply profile tumor and normal adjacent tissues from cancer patients to provide a more comprehensive proteomic and phosphoproteomic data than TCGA’s RPPA.

However, proteomics and phosphoproteomics abundance measurements of kinases do not reflect kinase activation state. To gain insights about potential kinase activity, kinase enrichment analysis (KEA) and other related methods have been developed. Briefly, kinase enrichment analysis accepts an input list of differentially phosphorylated proteins or phosphoproteins as putative substrates, and it then computes the overrepresentation of upstream kinases as possible global effectors of the observed changes in phosphorylation. There are several published kinase inference analysis applications that implement different methods of enrichment and network analysis to determine kinase activity from mass-spectrometry phosphoproteomics data. These computational methods utilize different underlying background kinase-substrate databases. For example, RoKAI is a method that considers the kinome network to determine the most likely areas in such a network that change in activity under different conditions [10]. A related computational approach was applied to phosphoproteomics data collected from acute myeloid leukemia (AML) cells to infer the most likely cell signaling pathways active within each cell line [11]. This analysis led to the inference of the best kinase inhibitors tailored for inhibiting the proliferation of each AML sample. By utilizing data from PhosphoSitePlus [12], IKAP implements a machine learning method to infer kinase activity from phosphoproteomics data [13]. IKAP was applied to infer kinase activity for over 100 kinases from several published datasets. Similarly, knowledge-based CLUster Evaluation (CLUE) [14] also utilized kinase-substrate knowledge from PhosphoSitePlus to infer kinase activity for time-series phosphoproteomics. Tools that implement more simple enrichment analysis methods include KSEA [15], KinasePA [16], KEA [17], and KEA3 [18]. Because phosphoproteomics data is noisy, partial, and our knowledge of kinase-substrate interactions is limited, the comprehensiveness of the datasets driving kinase inference from phosphoproteomics methods is critical. The recently published KEA3 platform [18] provides an improved kinase enrichment analysis method by leveraging a background database of integrated kinase-substrate interaction data assembled from over 20 resources which also include PhosphoSitePlus. KEA3 background databases encompass kinase-substrate interactions (KSIs), kinase-protein interactions (KPIs), and interactions derived from co-expression [19] and co-occurrence data [20].

In this work, leveraging hundreds of deep phosphoproteomic profiles from CPTAC studies, we comprehensively characterized tumor related kinase activities across five different cancer types using KEA3. Specifically, based on the deep phosphoproteomics data from paired tumor (T) and normal-adjacent-tumor (N) tissues from CPTAC studies, we derived patient specific T/N fold changes for each phosphosite and performed kinase enrichment analysis using subsets of phosphosites with either high or low T/N fold changes for each patient. The resulting kinase activity scores revealed key kinases differentiating tumor immune subtypes, which have implications for therapeutic approaches with small molecule inhibitors targeting specific protein kinases and kinome pathways. Importantly, we implemented the ProTrack Kinase Activity Portal (ProKAP), available from: https://pancan-kea3.cptac-data-view.org, to enable querying, visualizing, and downloading of the pan-cancer kinase activity scores, together with the corresponding proteomics and phosphoproteomics data, and tumor sample metadata. The ProKAP serves as a resource for hypothesis generation for cancer researchers.

## Methods

### Data summary

Publicly available proteomics and phosphoproteomics profiles were downloaded from the CPTAC data portal for the study cohorts of the following tumor types: clear cell renal cell carcinoma (ccRCC) [21], lung adenocarcinoma (LUAD) [22], uterine corpus endometrial carcinoma (UCEC) [23], colorectal cancer (CO) [24], and head and neck squamous cell carcinoma (HNSCC) [25]. Across these cohorts, there are 378 pairs of matching tumor (T) and normal adjacent tumor (N) samples with phosphosite abundance profiles. For each cancer type, we considered phosphosites detected in at least 50% of the samples. The total number of phosphosites across different cancer types range from ∼17K to ∼32K (Table 1).

**Table 1.** General stats of the 5 phosphoproteomics data sets from 5 CPTAC cancer studies. The second column provides the number of unique phosphosites identified in the preprocessed phosphoproteomics tables from each CTPAC cancer study. The other columns list the numbers of tumor samples, normal samples, and paired T/N samples in each cancer study.

| Cohort | Phosphosites | Tumor samples | Normal samples | Paired T-N samples |
| --- | --- | --- | --- | --- |
| CCRCC | 28,020 | 110 | 84 | 84 |
| COAD | 17,343 | 97 | 100 | 96 |
| HNSCC | 31,640 | 110 | 66 | 66 |
| LUAD | 32,792 | 115 | 108 | 102 |
| UCEC | 30,977 | 100 | 40 | 30 |

### Patient specific T/N phosphosite ratios

We first derived the (log2) T/N abundance ratios of all phosphosites for each patient by taking the differences between the log2 phosphosite abundances in the matched T and N samples. Based on these ratios, for each patient, we identified the top 500 proteins with the largest T/N phospho-abundance ratios. We refer to these lists as 500-up protein sets. Similarly, we identified the bottom 500 proteins with the smallest T/N phospho-abundance ratios for each patient and refer to these lists as 500-down protein sets. These protein sets were then inputted to the KEA3 Consensus Kinases Appyter to derive kinase activity rank scores.

### Kinase Enrichment Analysis

For each patient, KEA3 analysis was applied to the corresponding 500-up and 500-down protein sets separately. The 378×2 KEA3 analyses were carried out in batch-mode using the KEA3 API through the KEA3 Consensus Kinases Appyter available from the Appyters Catalog [26]. The Appyter can be accessed from: https://appyters.maayanlab.cloud/KEA3_Consensus_Kinases/ and is a part of the Appyter Catalog [26]. The output of the KEA3 Appyter is presented as a Jupyter Notebook [27] report. The report includes t-SNE [28] and UMAP [29] plots of the samples and kinases (colored by the tumor code). Tumor-specific kinases are highlighted in these plots. Interactive heatmaps visualized with clustergrammer [30] and stacked bar plots are also provided to visualize the scores of the top kinases for each sample. The Appyter instances for the analysis presented here are available via these permanent URLs. TNdiff2_top500:https://appyters.maayanlab.cloud/KEA3_Consensus_Kinases/823a777e835ce7926f55431ccb8552a0f1555875/ TNdiff2_bot500:https://appyters.maayanlab.cloud/KEA3_Consensus_Kinases/bd01544f4b418b76c617292d2476c6c350ad1b19/

We then focus on the KEA3 mean rank scores for the 529 kinases in each of the KEA3 runs. The lower rank scores indicate higher level of enrichment of kinase activities among the 500-up and 500-down protein sets, consequently suggesting increased or decreased kinase activation in T compared to N tissues. For each kinase, we then derived normalized KEA3 kinase activity scores by (1) Z-scoring the mean rank across all samples; and (2) flipping the sign. In this way, for each kinase, a larger score suggests a higher activity level. We further annotated the kinase activity scores based on the 500-up (or 500-down) protein sets as *Tumor-Up (or Tumor-Down) Kinases Activity scores*. The final kinase activity score matrices of 529 kinases for 378 cancer patients are hosted on the ProTrack Kinase Activity Portal (ProKAP).

### ProTrack Kinase Activity Portal (ProKAP) *Portal implementation*

ProKAP is a full-stack web application, architected with a Python Flask microframework backend and Vue.js reactive framework frontend. All data are stored in a JSON-based, NoSQL cloud database. The ProKAP source code is publicly available from GitHub at: https://github.com/WangLab-MSSM/protrack-pancan-kea3. The architectural diagram of the underlying software stack is shown in Fig. 1.

**Fig. 1.**
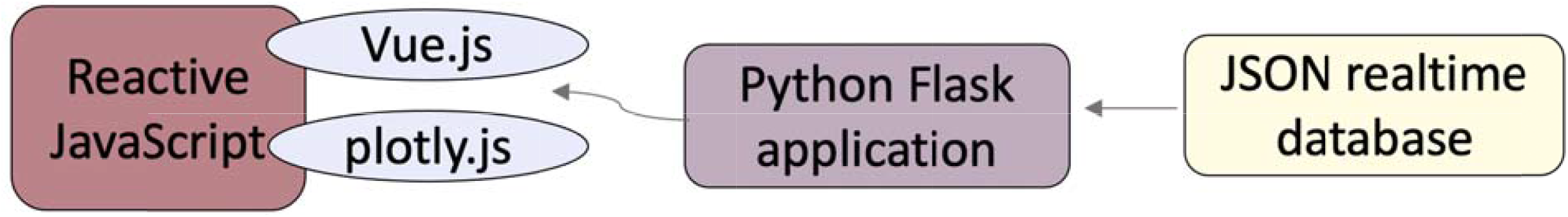
An architectural diagram of the underlying ProKAP software stack.

## Results

### Distribution of phosphosites T/N ratios

The distribution of the (log2) T/N ratio across all phosphosites and all patients of each tumor type appears to be balanced (Fig. 2). That is, for all the five tumor types, the log2-T/N-ratio distributions of all the phosphosites are close to symmetric about 0. The most up regulated phosphosite in tumors compared to normal tissues based on the T/N ratio was observed for the phosphosite KRT79-S20 (log2 T/N=8.389) in one of the HNSCC samples, suggesting potential disturbed cellular structural transformations in this HNSCC tumor. The top 5 most up- and down-regulated phosphosites based on their T/N ratios are provided in Table 2.

**Table 2.** Top 5 phosphosites that display the greatest change between paired T and N samples across all tumor types. The top (bottom) table lists the phosphosites that increased (decreased) in abundance in tumor compared to adjacent normal tissues.

| HGNC symbol | T/N fold change | Tumor Type | sample | GENECODE34 Site |
| --- | --- | --- | --- | --- |
| KRT79 | 8.3888395 | HNSCC | C3N-01758 | ENSP00000328358.5 |
| IGF2BP3 | 8.215243 | LUAD | C3N-02379 | ENSP00000258729.3 |
| KRT75 | 8.07009418 | HNSCC | C3N-03928 | ENSP00000252245.5 |
| MAP2 | 8.02295 | CCRCC | C3N-00492 | ENSP00000353508.4 |
| SQSTM1 | 8.0194702 | CCRCC | C3L-00360 | ENSP00000374455.4 |

| Tumor Type | sample | T/N fold change | HGNC symbol | GENECODE34 Site |
| --- | --- | --- | --- | --- |
| HNSCC | C3N-01947 | -9.960516 | KRT76 | ENSP00000330101.2 |
| HNSCC | C3L-02651 | -9.610841 | MYH1 | ENSP00000226207.5 |
| HNSCC | C3N-03028 | -9.300213 | KRT76 | ENSP00000330101.2 |
| HNSCC | C3N-04279 | -9.141568 | CRNN | ENSP00000271835.3 |
| HNSCC | C3N-04279 | -9.061656 | CRNN | ENSP00000271835.3 |

**Fig. 2.**
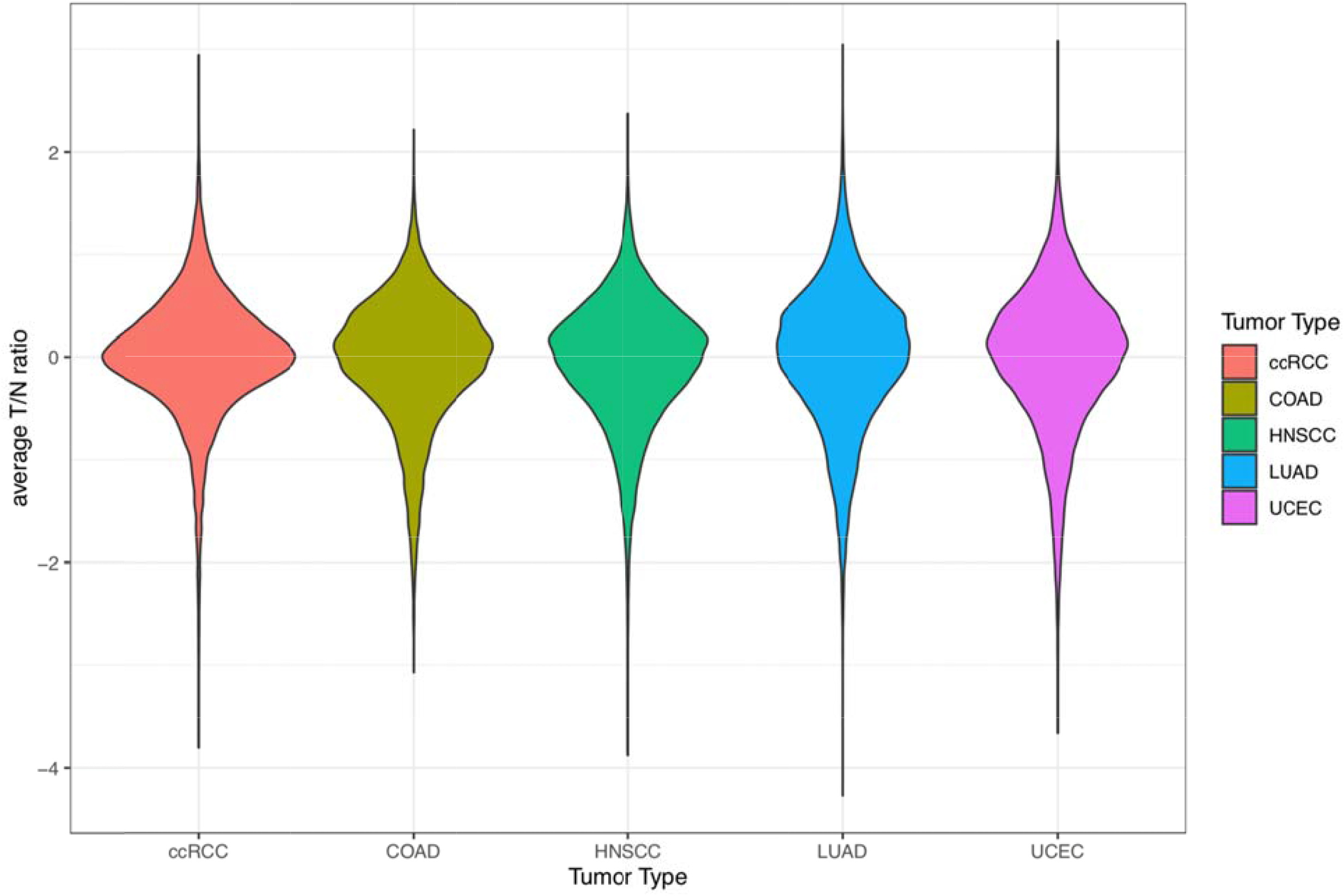
Distributions of the mean tumor-normal ratios of phosphosites for each tumor type.

### Derived KEA3 Kinase Activity Scores and the ProTrack Kinase Activity Portal (ProKAP)

For each patient, the top 500 proteins that had their phosphosites altered in tumor tissues, either up- or down-regulated, were submitted to KEA3 for kinase enrichment analysis. A t-SNE plot that displays the patient similarity based on their 529 kinase activity scores can distinctly separate tumor samples by the different cancer subtypes, suggesting that the same cancer subtypes are preserved by the specific kinase activity scores (Fig. 3).

**Fig. 3.**
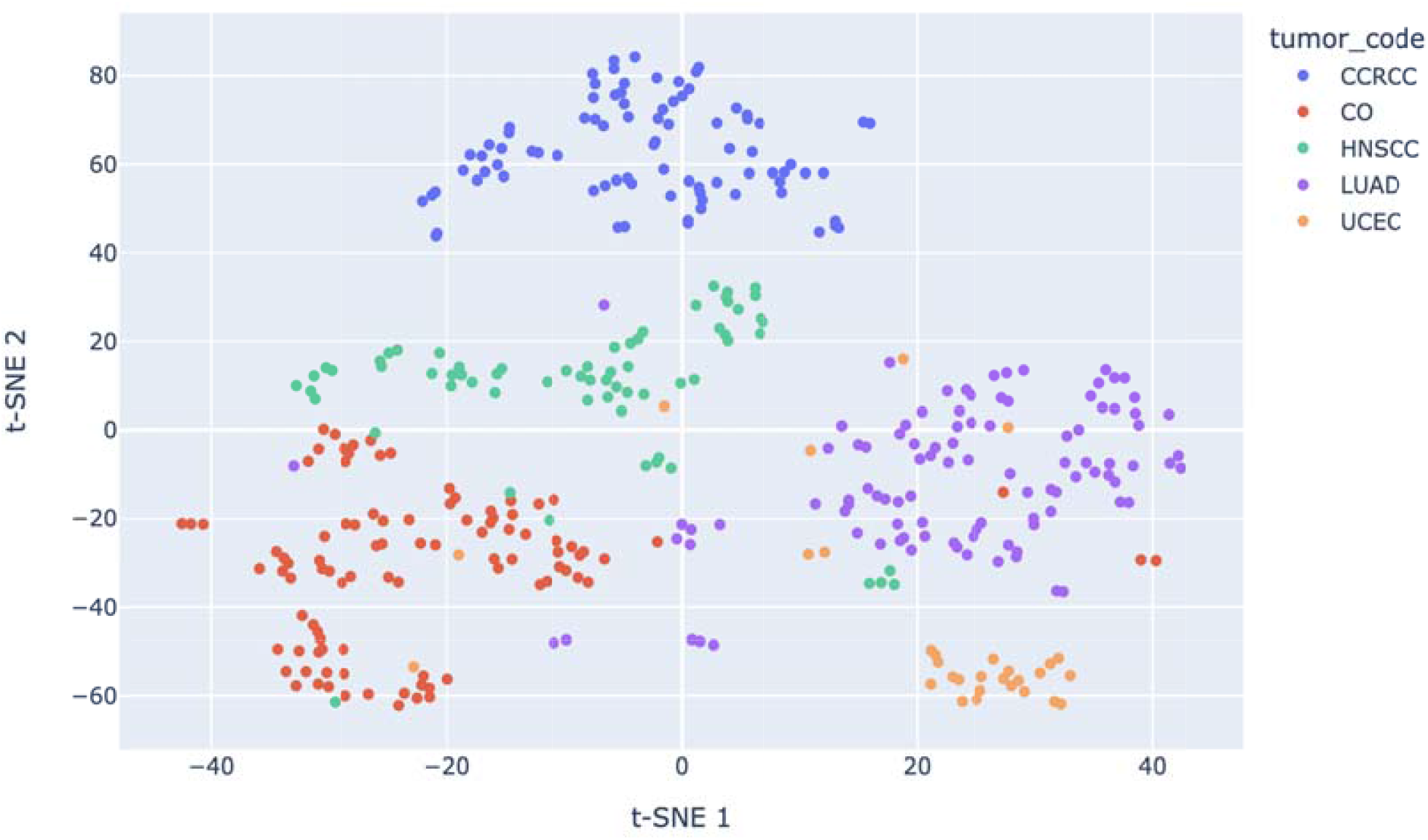
t-SNE plot of 378 tumor samples based on KEA3 mean rank scores (kinase enrichment vectors). Colors of the dots represent the tumor types.

The derived KEA3 kinase activity scores for 529 kinases in 378 tumors of 5 different cancer types can be queried, visualized, and downloaded through ProKAP at https://pancan-kea3.cptac-data-view.org. The portal provides an input form element that can be used to submit a set of kinases and then multiple interactive heat maps are generated. The heat maps provide visual comparison of different assessments of kinase activity scores across multiple cancer types. Once the list of kinases is submitted, the users can access a “Customize and Download” panel. From this panel, the user can perform a cohort query selection, by filtering the results based on tumor type and clinical features, including stage, grade, and survival status. The entire heat map can then be sorted by any given molecular or clinical track. This enables automatically aligning all data types based on a single sorted data type and visually detecting patterns between and across kinases. For a given inputted kinase, the heatmap also includes protein abundance fold change and phosphosite abundance for purported substrates. These additional tracks can be used to visually confirm the alignment of the kinase score with the differential protein abundance of a given kinase, as well as explore the kinase-phosphosite relationships across cancers (Fig. 4).

**Fig. 4.**
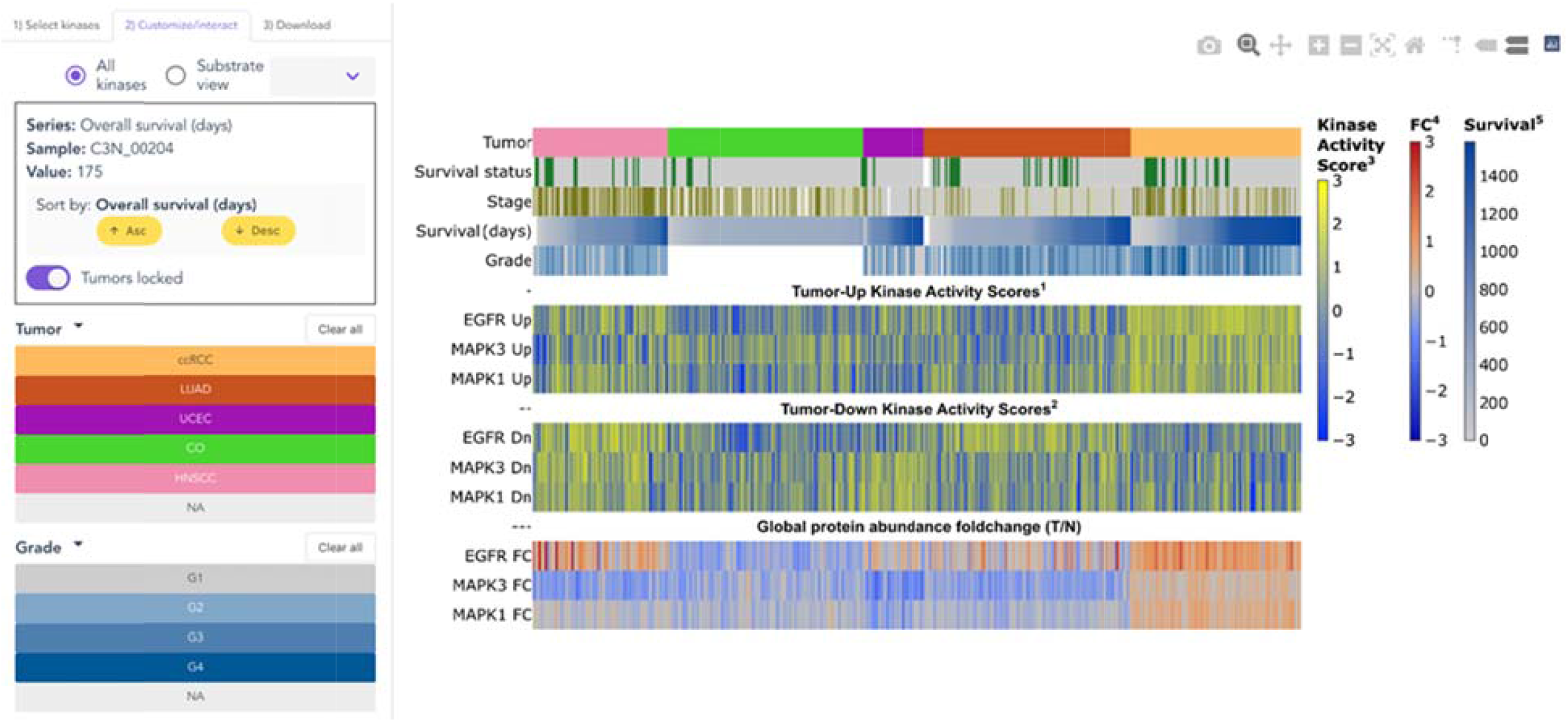
The ProKAP interface. Given an input list of kinases, the heatmap displays Tumor-Up and Tumor-Down kinase activity scores across 378 cancer patients. Features of the interface include a substrate view where users can visualize the relationship between kinase enrichment score for a given kinase and the corresponding fold change of expression in tumor versus normal tissue of its purported phospho-substrates. Users can click on any series then sort the entire heatmap in ascending or descending order. This enable exploring trends. Users can also filter cohorts by various clinical features, including tumor type, grade, stage, and survival status.

### Kinase activity scores differentiate key kinases in ccRCC immune subtypes

To illustrate how kinase activity scores can be used to illuminate biological mechanisms in these tumors, we investigated highly ranked kinases relating to immune activation and evasion in ccRCC. The CPTAC ccRCC study [21] identified four immune subtypes based on cell type composition of tumor tissues inferred through deconvolution analysis of RNA-seq data. These four immune subtypes represent distinct molecular and microenvironment signatures and were annotated as: CD8+ inflamed, CD8-inflamed, VEGF, and metabolic immune desert subtypes. Interestingly, the KEA3 analysis applied to the phosphoproteomics data provides additional insights on the mechanistic explanation of the observed molecular differences across immune subtypes at the cell signaling level. Specifically, to gain insights on immune signaling pathways that have therapeutic implications among each immune subtype, we derived kinase signatures specific to each subtype by testing the association between kinase activity scores and the subtype labels. We focused on the *Tumor-Up kinase activities scores* (derived based on the 500-up gene lists) and compared these scores in one subtype of ccRCC tumors with the rest using two sample t-tests. Kinases with significantly higher activity scores for each immune subtype are shown in Fig. 5A. First, as a positive control, we confirmed that PRKCQ, a positive effector of T-cell receptor signaling and functioning mainly in T lymphocytes [31], has the highest kinase activity scores in CD8+ inflamed ccRCC tumors, which are a subset of tumors characterized by a high level of CD8+ T-cell infiltration (left panel, Fig. 5B). However, upregulated activity of PRKCQ in CD8+ inflamed tumors cannot be revealed based on its global abundance (right panel, Fig. 5B). It’s worth noting that, PRKCQ is one of the “dark kinases”, a list of understudied kinases with limited information about their molecular function and phosphosite targets [32]. Thus, our observation that PRKCQ is a top ranked kinase in this subtype demonstrates the effectiveness of the pipeline to characterize kinase activity based on phosphoproteomics data through leveraging KEA3. Moreover, there is a growing body of research that is illuminating the role of PRKCQ in cancer initiation and progression [33]. For example, it was reported that PRKCQ inhibition enhances chemosensitivity of triple-negative breast cancer, through regulation of Bim expression [34]. Our findings further suggest that PRKCQ could be studied as a therapeutic target of interest for an immune-specific subtype of ccRCC tumors.

**Fig. 5.**
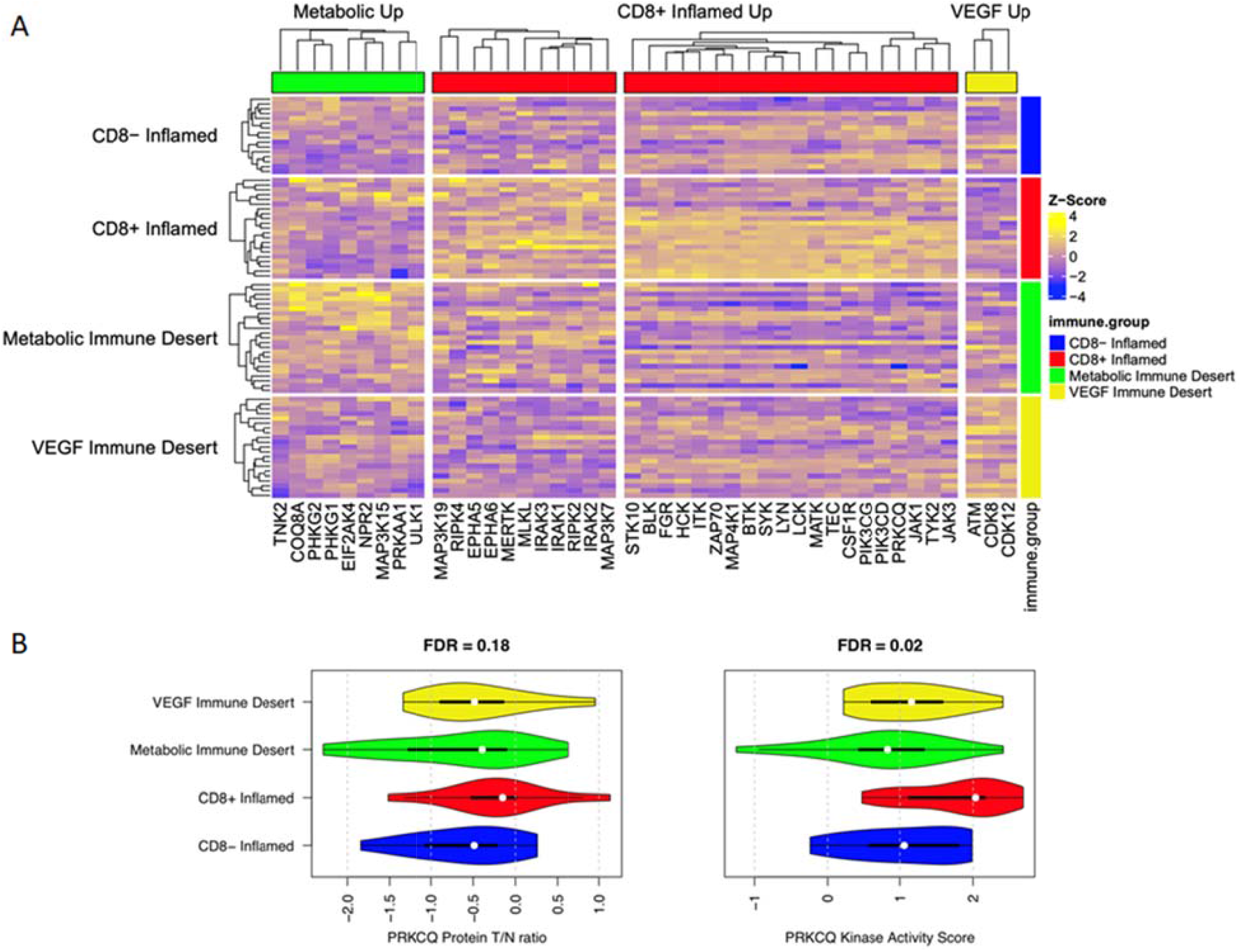
Kinase activity scores associated with immune subtypes in ccRCC. A) Heatmap of normalized kinases activity scores for kinases with upregulated activation in each ccRCC immune subtype. B) Distribution of the tumor/normal differences (left) and the corresponding kinase scores (right) of PRKCQ across different immune subtypes.

In addition to PRKCQ, bruton tyrosine kinase (BTK) is another interesting kinase that showed strikingly elevated kinase activities in CD8+ Inflamed tumors (Fig. 5B). BTK is a critical kinase that interconnects BCR signaling, Toll-like receptor (TLR) signaling, and chemokine receptor signaling. There is a BTK inhibitor that is emerging as a promising novel agent displaying potential efficiency in B-cell malignancies. This inhibitor has been applied to many other cancer types in more than 70 trials in the past 3 years [35]. Moreover, CD8+ inflamed tumors are marked by an inferred increase in activity of the innate immune kinases: IRAK4, IRAK3, IRAK2, and notably IRAK1 -- a reported therapeutic target for breast cancer [36]. These observations should be considered as novel hypotheses of personalized drug selections for individuals or subsets of patients. As to other ccRCC immune subtypes, the VEGF Immune Desert tumors suggest activation of ATM and cyclin-dependent kinases CDK8 and CDK12. This has implications for current approaches in the suppression of angiogenesis in tumors, by targeting cyclin-dependent kinases [37]. Metabolic Immune Desert tumors were enriched in activated kinases PHKG1, PHKG2, TNK2, NPR2, COQ8A, PRPKAA1, ULK1, EIF24K4, and MAP3K15.Overall, these observations are consistent with observations made with RNA-seq data and clinical characterization of tumor subtypes of ccRCC.

## Discussion

Protein kinases are common targets for inhibition by anticancer agents. To facilitate the discovery of potential kinase targets in personalized medicine, here we derived kinase activity scores for 378 tumors extracted from patients of five cancer types profiled by deep phosphoproteomics from CPTAC. We utilized KEA3 to perform kinase enrichment analysis using gene sets whose phosphosite abundances were up- or down-regulated in the tumors compared with matched adjacent normal tissues from each patient. The inferred kinase activity scores are hosted on the ProTrack Kinase Activity Score Portal (ProKAP) for the community to glean more knowledge from this useful dataset. Users of ProKAP can query, visualize, and download the kinase activity scores from this web portal. By exploring kinase enrichment scores across a diverse group of cancers using ProKAP, researchers can investigate whether their targets of interest are applicable to individual patients, cancer subtypes, and cancer types. We illustrate the usage of the kinase activity scores by identifying kinases whose increased activities were associated with different immune subtypes in ccRCC. For example, over the past several years, one growing family of kinase inhibitors target the bruton tyrosine kinase (BTK), particularly for treating chronic lymphocytic leukemia (CLL) and mantle cell lymphoma (MCL). While BTK inhibitors primarily target B cell malignancies, there are recent clinical studies applying these drugs to a wider range of other cancer types [35]. However, response to BTK inhibitors in kidney cancer is inconsistent, ranging from no response to a reported 22-month clinical response [38]. Stratification of patients by immune subtype may facilitate more targeted clinical studies. BTK demonstrated increased enrichment in the CD8+ Inflamed subtype (Fig. 5), suggesting that tumors of this subtype may be more responsive to BTK inhibition. The results from our analysis shed light on immune signaling pathways that have therapeutic implications among each immune subtype.

While here we provide analysis of five cancer types, we are aware that more samples from more cancer types are actively profiled by CPTAC with phosphoproteomics. We plan to analyze newly released data sets and expand the ProKAP resource on a regular basis. In addition, integration with other omics datasets such as RNA-seq and scRNA-seq is expected. For example, by analyzing matching RNA-seq with the tool ChEA3 [39] may enable us to link kinase activity scores with transcription factor activity scores [40]. There are some limitations with this current work. First, the databases driving kinase enrichment analysis in KEA3 may be affected by biases toward over-reporting well-studied kinases. Additionally, KEA3 is applied at the protein level, missing important nuances that occur at the phosphosite level. Future research may include the application of kinase enrichment to phosphosites directly, including single-cell considerations, and spatial proteomics and transcriptomics data.

## Acknowledgements

This work is partially supported by NIH grants U24CA210993 to PW and U24CA224260, U54HL127624, and OT2OD030160 to AM.

## Notes

### Competing Interest Statement

The authors have declared no competing interest.

